# zFACE: <u>F</u>acial <u>A</u>nalytics from a <u>C</u>oordinate <u>E</u>xtrapolation System for Developing Zebrafish

**DOI:** 10.1101/2022.07.26.501188

**Authors:** Lorena Maili, Oscar E. Ruiz, Philip Kahan, Stephen T. Larson, S. Shahrukh Hashmi, Jacqueline T. Hecht, George T. Eisenhoffer

## Abstract

Facial development requires a complex and coordinated series of cellular events, that when perturbed, can lead to structural birth defects. A standardized quantitative approach to quickly assess morphological changes could address how genetic or environmental inputs lead to differences in facial development. Here we report on a method to rapidly analyze craniofacial development in zebrafish embryos that combines a simple staining and mounting paradigm with <u>F</u>acial <u>A</u>nalytics based on a <u>C</u>oordinate <u>E</u>xtrapolation system, termed zFACE. Confocal imaging of frontal/rostral mounted embryos generates high-resolution images to capture facial structures and morphometric data is quantified based on a coordinate system that assesses 26 anatomical landmarks present at defined times in development. The semi-automated analysis can be applied to embryos at different stages of development and quantitative morphometric data can detect subtle phenotypic variation. Shape analysis can also be performed with the coordinate data to inform on global changes in facial morphology. We applied this new approach to show that loss of *smarca4a* in developing zebrafish leads to craniofacial anomalies, microcephaly and alterations in brain morphology. These changes are characteristic of humans with Coffin-Siris syndrome (CSS), a rare genetic disorder associated with mutations in *SMARCA4* that is defined by anomalies in head size, intellectual disabilities and craniofacial abnormalities. We observed that *smarca4a* is expressed in craniofacial tissues and our multivariate analysis facilitated the classification of *smarca4a* mutants based on changes in specific phenotypic characteristics. Together, our approach provides a way to rapidly and quantitatively assess the impact of genetic alterations on craniofacial development in zebrafish.

## INTRODUCTION

Vertebrate craniofacial development requires the complex orchestration of cellular processes, molecular signals and interactions between different cell types [1,2]. During the course of organ development, interactions between different tissues are critical in the regulation of cellular proliferation, migration, and apoptosis to develop the intricate features that comprise the face [2,3]. Failure to efficiently coordinate the specification, migration, proliferation and apoptosis of cells can lead to craniofacial malformations and structural birth defects, including cleft lip and/or palate, craniosynostosis and facial dysostosis [4,5]. Many developmental syndromes, such as Stickler, Van der Woude and Coffin-Siris Syndromes (CSS), include craniofacial anomalies in addition to other congenital malformations [4,6]. One impediment to understanding how genetic variants promote craniofacial anomalies has been our ability to visualize the complex and coordinated cellular interactions during craniofacial development *in vivo*.

Animal models, such as mouse, chicken, frog and zebrafish, provide an important avenue to gain better mechanistic and genetic insights into human craniofacial development [5]. Zebrafish offer many advantages as a model system that make them ideal for detailed craniofacial studies — they develop externally and generate large numbers of transparent embryos, which permits unparalleled high-resolution imaging of specific cell types and structures in living vertebrates [7,8,9,10,11,12]. Zebrafish craniofacial development, particularly craniofacial bone and cartilage formation, has been well characterized and is comparable to aminote craniofacial development. For example, the anterior neurocranium/ethmoid plate is thought to be functionally analogous to palate development in mammals [13,14]. Zebrafish have been used to systematically test the function of genes associated with birth defects in humans [15,16,17,18,19,20,21,22,23,24,25], and an array of conserved genetic variants/mutations already exist that display altered craniofacial development [26,27]. Yet, a standardized, unbiased method that can be used to quantitatively assess phenotypic changes in craniofacial structures resulting from these mutations is not currently available.

Geometric morphometrics (GMM) is a quantitative method used to measure and statistically test for variation(s) in shape [28]. GMM methods have been applied to skeletal and soft tissue data to evaluate craniofacial morphology in the clinical setting and to better understand human craniofacial disorders, such as cleft lip and palate, craniosynostosis, ectodermal dysplasia, and neurodevelopmental disorders [29,30,31,32,33,34,35]. The presentation of craniofacial abnormalities and syndromes often display phenotypic heterogeneity, and the use of GMM provides an important approach that captures critical information that would otherwise be missed using only a discrete categorical variable such as the presence or absence of a normal or abnormal phenotype [28]. For example, GMM has been used to identify facial differences in unaffected relatives of individuals with non-syndromic cleft lip and palate and strain-specific differences in embryonic facial shape underlying susceptibility to developing orofacial clefts in mice [32,36].

To leverage the power of the GMM approach for zebrafish embryos, we have developed a novel and easily implemented method, termed zFACE (<u>F</u>acial <u>A</u>nalytics based on a <u>C</u>oordinate <u>E</u>xtrapolation system), to visualize the developing rostral/frontal craniofacial region and analyze quantitative facial morphometric data in a semi-automated way. We apply zFACE to show that loss of *smarca4a* modifies craniofacial morphology in zebrafish embryos and identify regional differences that contribute to the observed altered facial dimensions. These results support *smarca4a* mutant zebrafish as an animal model to provide insight into the craniofacial abnormalities associated with Coffin-Siris syndrome (CSS), a rare genetic disorder associated with mutations in *SMARCA4/BRG1*. Together, our GMM-based approach and its application demonstrate the power of zFACE to extend our understanding of phenotypic changes in craniofacial development associated with genetic alterations in a vertebrate embryo.

## RESULTS

### zFACE – <u>F</u>acial <u>A</u>nalytics from a <u>C</u>oordinate <u>E</u>xtrapolation system

Zebrafish embryos develop externally and provide a unique opportunity to visualize development [7,8,18]. While zebrafish craniofacial development has been well characterized, most studies have used lateral and ventral views of the developing face. To visualize additional craniofacial anatomical structures, we first established a simple imaging paradigm using DAPI, a stain which labels the nucleus of every cell, along with face-on rostral mounting method using low-melt agarose (**Figure 1A**). This method can be easily adapted to accommodate different developmental stages, when the orientation of the face with respect to the rest of the body varies, by simply adjusting the mounting angle of the specimen. The resulting images capture structures such as the neuromasts, olfactory placodes, eyes, oral cavity, and frontonasal, maxillary and mandibular regions (**Figure 1B**). Thus, this rostral mounting paradigm provides high-resolution images that reveal important facial information compared to the commonly used lateral and ventral views (**Figure 1A**).

**Figure 1.**
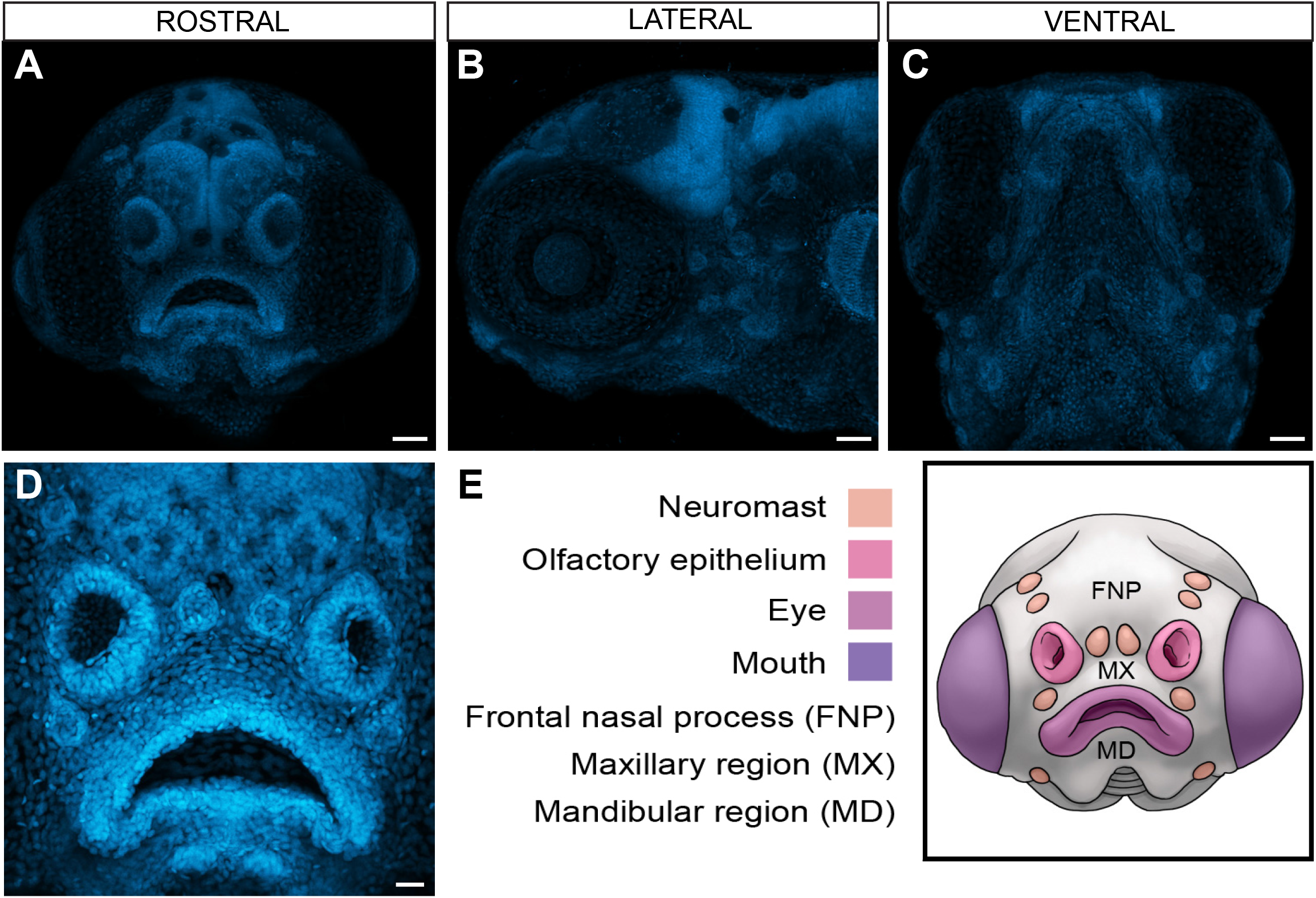
Craniofacial features visualized in frontal/rostral mounts by DAPI stain and confocal microscopy. **A-D.** Rostral (**A,D**), lateral (**B**) and ventral (**C**) views of developing zebrafish larvae at 5 days post fertilization (dpf). **E.** Schematic representing the facial anatomical structures visualized in the rostral view, including neuromasts, olfactory placodes, eyes, mouth, frontonasal region (FNP) and maxillary and mandibular regions (MX and MD). Scale bars in A-C = 50μm, D = 20μm.

We applied geometric morphometrics (GMM) to quantitatively analyze facial form based on this newfound ability to identify facial features and landmarks using zFACE. Based on the information captured by the confocal images, 26 easily-identifiable landmarks during the embryonic and early larval stages of development were established (**Figure 2 and Supplemental Table 1**). Given the complex shape of the oral cavity, and that many craniofacial abnormalities affect the mouth, approximately one-third of the landmarks were assigned to this area to provide better resolution of changes in morphology for this region. Additionally, landmarks were chosen and named to parallel those used in human GMM studies [32,37]. Using these established landmarks, an automated calculation of 39 different linear distances, angles, and areas was generated to determine the localization of landmarks relative to each other. Additional statistical approaches were then applied to evaluate overall facial shape (**Figure 2 and Supplemental Table 1**). Together, these analyses can be used to quantify phenotypic information resulting from genetic perturbation or environmental exposures and provide valuable information about craniofacial form.

**Figure 2.**
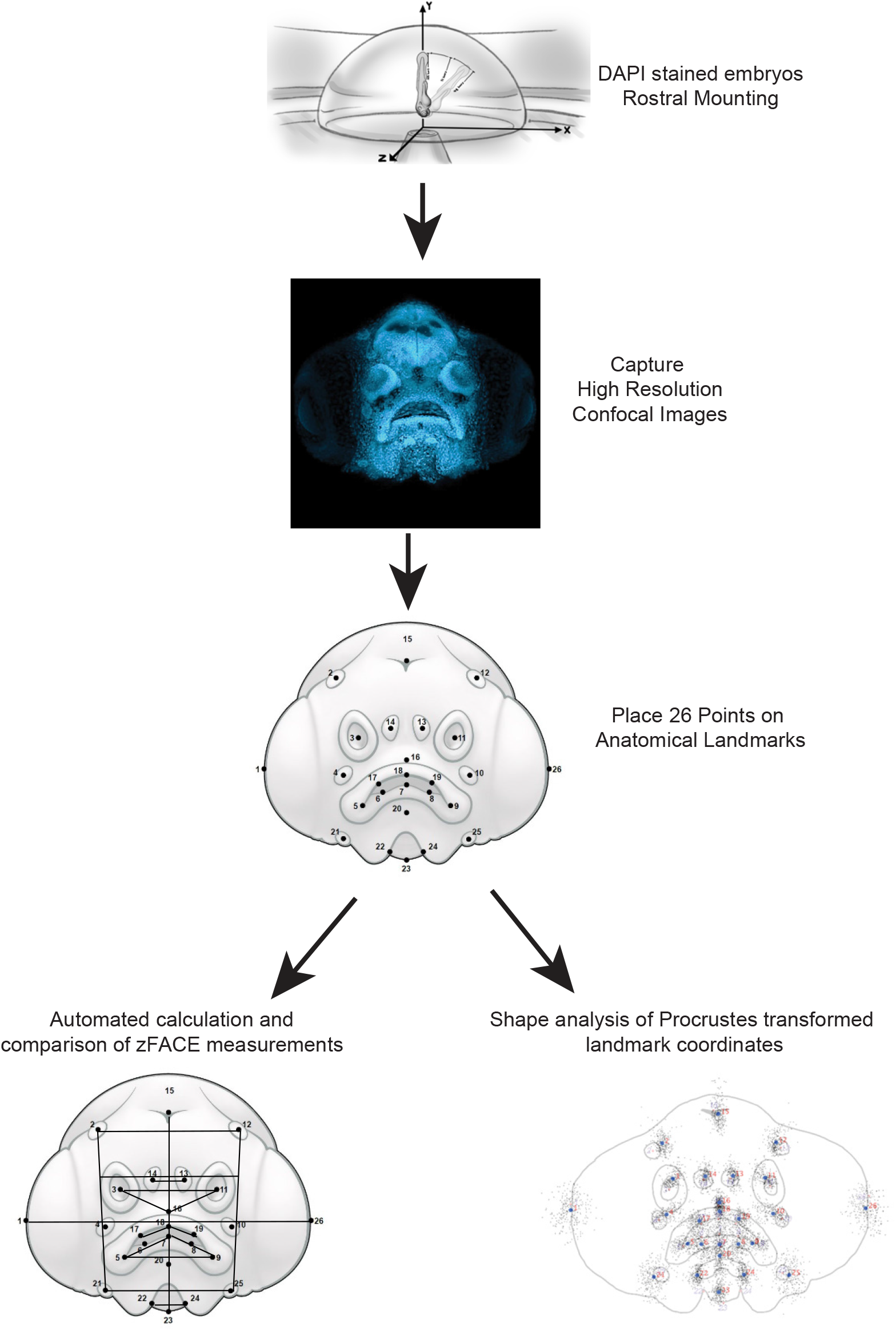
zFACE workflow for morphometric analysis of the developing zebrafish craniofacial region. Schematic representing the steps in zFACE analysis, where larvae are mounted and images are then acquired; points are then placed on the 26 anatomical landmarks; and either feature-based zFACE calculations or overall shape analyses can be performed.

### Morphometric analysis of facial development in zebrafish

To assess the ability to detect changes in specific anatomical structures and locations, we used zFACE to investigate facial development over time. Images of rostrally mounted zebrafish embryos from 48 hours post-fertilization (hpf) to 6 days post-fertilization (dpf) were acquired by confocal microscopy and compared to scanning electron microscope (SEM) images. The morphology of the soft tissue and anatomical structures was similar in both conditions, indicating that fixation and the DAPI stain, mounting technique or confocal capture does not induce artifacts in facial form (**Figure 3A**). The resulting confocal and SEM images revealed the emergence of specific anatomical structures and regional changes that occurred at defined times throughout development. For example, between 2 and 3 dpf, the oral cavity expands into a semi-circle morphology, while the olfactory placodes shift ventrally. Between 3 and 4 dpf, the biggest changes include a narrowing of the face and an enlargement of the mouth, while between 4 and 5 dpf, the midface widens and the oral cavity becomes crescent shaped. Comparison of 5 and 6dpf confocal images did not discern any major changes in morphology. All of the 26 anatomical landmarks used by zFACE were not present until 3 dpf, and therefore, quantitative analysis included the course of development from 3 to 6 dpf.

**Figure 3.**
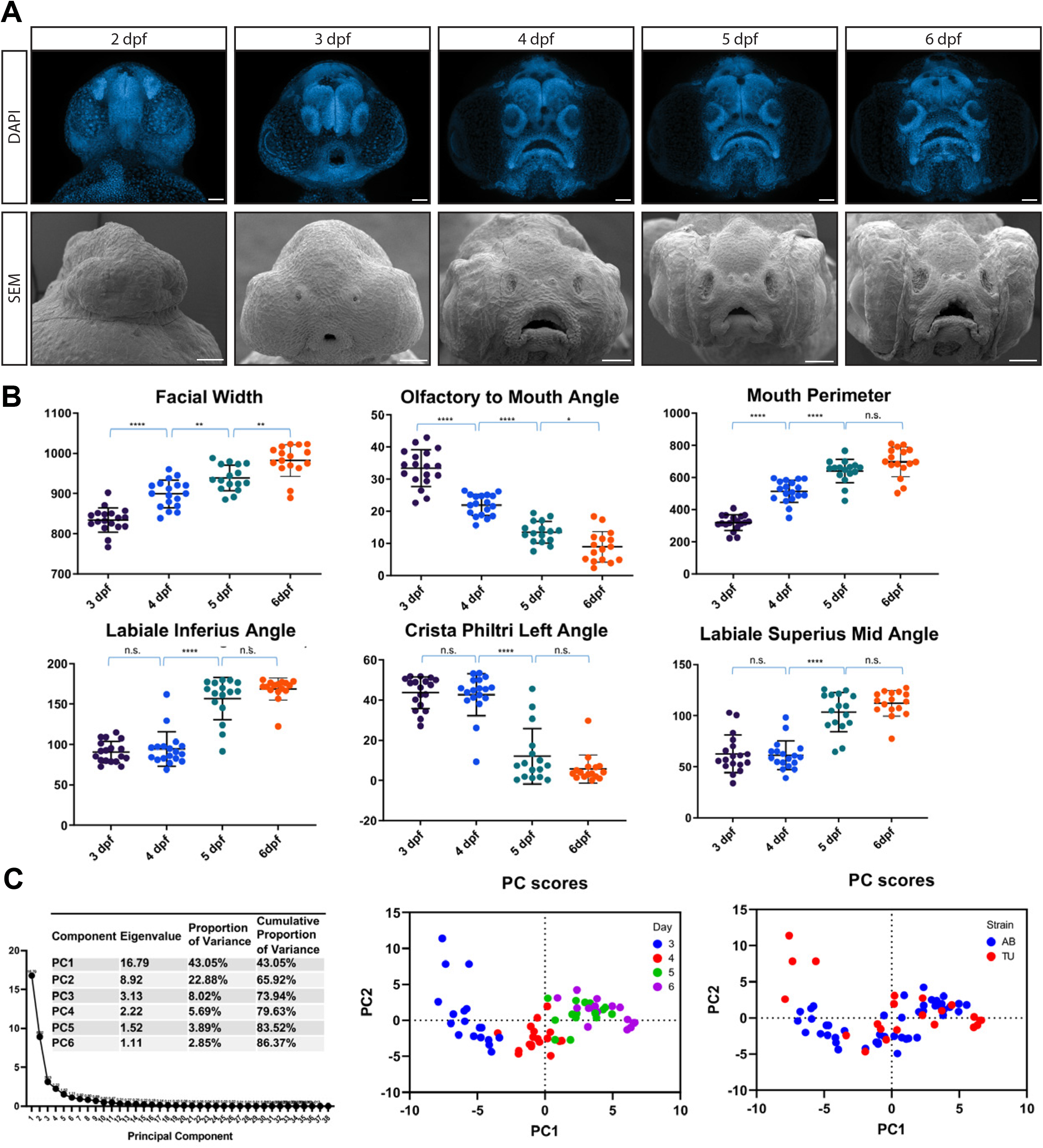
Time series of zebrafish craniofacial development. **A.** Zebrafish larvae were analyzed from 2 - 6 days post fertilization (dpf). Scanning electron microscope (SEM) images confirmed the soft tissue morphology captured by DAPI staining and confocal microscopy used in zFACE. **B.** Changes in zFACE features were followed across developmental time points and alterations in 27 measurements were determined (representative changes shown in the top and bottom graph panels). Specifically, 21 zFACE measurements changed between 3 and 4 dpf, 18 changed between 4 and 5 dpf, and 6 changed between 5 and 6 dpf. **C**. Untransformed zFACE features were subjected to multivariate statistical testing using principal component analysis (PCA). The first 2 components captured 66% of the variance in the dataset. There were differences in developmental times that were captured across PC1, with no strain-specific differences observed. Scale bars = 50μm.

The resulting zFACE measurements were compared across developmental time points and strains. Nineteen measurements, including facial width and height, mouth width and height, olfactory to mouth angles, as well as multiple neuromast measurements, were observed to be significantly different between 3 and 4 dpf (**Table 1, Figure 3B**). While the oral cavity angles did not differ between these early developmental timepoints, comparison of 4 and 5 dpf revealed 12 significantly different measurements, which included mouth width and height as well as labiale inferius, crista philtri, labiale superius and labiale inferius angles (**Table 1, Figure 3B**). Additionally, when later-stage (5 dpf and 6 dpf) larvae were compared, only 6 measurements were different, suggesting that facial morphology was very similar between these two time points (**Table 1**, **Figure 3B**). Interestingly, 9 measurements showed a daily increase, 8 showed a daily decrease, while 10 showed a non-linear change (**Table 1**). Comparison of facial development between the two most commonly used laboratory zebrafish strains, AB and TU showed relatively few significant differences (**Supplemental Table 2**) among the 39 different measurements, suggesting that the strains are very similar in both the timing of anatomical changes and overall facial morphology [38]. This analysis identified time points in developing zebrafish when facial dimensions are most dynamic and ones where face shape remains more stable.

**Table 1.** zFACE results showing phenotypic differences in craniofacial morphology between developmental days. Bolded values are significant, italicized values meeting 0.05 cutoff are also shown.

| Trend | zFACE Measurement | Trend | ANOVA | 3 dpf vs. |  |  | 4 dpf vs |  | 5 dpf vs |
| --- | --- | --- | --- | --- | --- | --- | --- | --- | --- |
|  |  |  |  | 4 dpf | 5 dpf | 6 dpf | 5 dpf | 6 dpf | 6 dpf |
| Daily increase | Width |  | <0.0001 | <0.0001 | <0.0001 | <0.0001 | 0.006 | <0.0001 | 0.003 |
|  | Height |  | <0.0001 | <0.0001 | 0.001 | 0.0006 | -- | -- | -- |
|  | Mouth Width |  | <0.0001 | <0.0001 | <0.0001 | <0.0001 | <0.0001 | <0.0001 | -- |
|  | Olfactory to Mouth 3 |  | <0.0001 | <0.0001 | <0.0001 | <0.0001 | <0.0001 | <0.0001 | 0.02 |
|  | Mouth to Chin |  | <0.0001 | <0.0001 | <0.0001 | <0.0001 | -- | -- | -- |
|  | Neuromast Width |  | <0.0001 | 0.01 | 0.003 | <0.0001 | -- | -- | -- |
|  | Area Bottom |  | <0.0001 | <0.0001 | <0.0001 | <0.0001 | -- | 0.001 | -- |
|  | Area Combined |  | <0.0001 | <0.0001 | <0.0001 | <0.0001 | -- | -- | -- |
|  | Mouth Perimeter |  | <0.0001 | <0.0001 | <0.0001 | <0.0001 | <0.0001 | <0.0001 | -- |
| Increase at 4dpf | Labiale Inferius Angle |  | <0.0001 | -- | <0.0001 | <0.0001 | <0.0001 | <0.0001 | -- |
|  | Labiale Superius Mid Angle |  | <0.0001 | -- | <0.0001 | <0.0001 | <0.0001 | <0.0001 | -- |
| Increase then decrease | Mouth Area |  | <0.0001 | <0.0001 | 0.006 | 0.0009 | 0.01 | -- | -- |
|  | Mouth Height |  | <0.0001 | <0.0001 | -- | -- | <0.0001 | <0.0001 | -- |
|  | Chin Width |  | <0.0001 | <0.0001 | <0.0001 | 0.0005 | -- | -- | -- |
| Daily decrease | Olfactory to Mouth |  | <0.0001 | <0.0001 | <0.0001 | <0.0001 | <0.0001 | <0.0001 | 0.04 |
|  | Olfactory to Mouth 2 |  | <0.0001 | <0.0001 | <0.0001 | <0.0001 | <0.0001 | <0.0001 | 0.02 |
|  | Neuromast Angle 1 |  | <0.0001 | 0.006 | <0.0001 | <0.0001 | 0.02 | <0.0001 | 0.002 |
|  | Neuromast Angle 2 |  | <0.0001 | <0.0001 | <0.0001 | <0.0001 | 0.01 | <0.0001 | 0.01 |
|  | Neuromast Height |  | <0.0001 | <0.0001 | <0.0001 | <0.0001 | 0.006 | 0.0004 | -- |
|  | Mid Neuromast Width |  | <0.0001 | <0.0001 | <0.0001 | <0.0001 | -- | -- | -- |
|  | Average Length Olfactory to mouth |  | <0.0001 | <0.0001 | <0.0001 | <0.0001 | -- | -- | -- |
|  | Area Top |  | <0.0001 | <0.0001 | <0.0001 | <0.0001 | 0.005 | 0.0004 | -- |
| Decrease after 4dpf | Crista Philtri Left Angle |  | <0.0001 | -- | <0.0001 | <0.0001 | <0.0001 | <0.0001 | -- |
|  | Crista Philtri Right Angle |  | <0.0001 | -- | <0.0001 | <0.0001 | <0.0001 | <0.0001 | -- |
|  | Labiale Inferius Left Angle |  | <0.0001 | -- | <0.0001 | <0.0001 | <0.0001 | <0.0001 | -- |
|  | Labiale Inferius Right Angle |  | <0.0001 | -- | <0.0001 | <0.0001 | <0.0001 | <0.0001 | -- |
|  | Labiale Inferius Diff |  | 0.0003 | 0.01 | 0.02 | 0.0002 | -- | -- | -- |
| No significant change | Olfactory Distance |  | 0.005 | -- | -- | -- | -- | 0.009 | 0.03 |
|  | Mid Olfactory to Chin height |  | 0.0013 | 0.0008 | -- | 0.02 | -- | -- | -- |
|  | Olfactory to Mouth Difference |  | 0.002 | 0.02 | 0.01 | 0.002 | -- | -- | -- |
|  | Alternate Height |  | 0.007 | 0.003 | -- | -- | -- | -- | -- |
|  | Crista Philtri Diff |  | 0.001 | -- | 0.01 | 0.003 | -- | -- | -- |
|  | Upper Lip Width |  | -- | -- | -- | -- | -- | -- | -- |
|  | Lower Lip Width |  | -- | -- | -- | -- | -- | -- | -- |
|  | Neuromast Angle Difference |  | -- | -- | -- | -- | -- | -- | -- |
|  | Libiale Superius Angle |  | -- | -- | -- | -- | -- | -- | -- |
|  | Chelion Left Angle |  | -- | -- | -- | -- | -- | -- | -- |
|  | Chelion Right Angle |  | -- | -- | -- | -- | -- | -- | -- |
|  | Chelion Diff |  | -- | -- | -- | -- | -- | -- | -- |

To understand which zFACE measurements contribute the most variance throughout development, we performed multivariate principal component analysis (PCA) of the combined AB and TU groups for all timepoints. Six principal components met the Kaiser cutoff and collectively accounted for 86% of the variation in the dataset (**Figure 3C**). Examination of the principal component (PC) loadings were then used to identify the measurements driving variance along each component. Results showed that neuromast width significantly loaded into PC1, while mouth height, mouth area, labiale inferius angle and left and right crista philtri angles significantly loaded into PC2. The labiale superius and left and right chelion angles significantly loaded into PC3 (**Supplemental Table 2**). These data suggested that measurements associated with the morphology of the mouth significantly contributed to the differences between developmental stages. Further, examination of PC plots for the first two components revealed clustering of groups by developmental day along PC1 (**Figure 3C**), while no strain-specific clusters were observed. These results indicate that unbiased analyses can also be used to understand how facial morphology changes during early craniofacial development, and support the results obtained from the automated zFACE calculations.

We next applied shape analysis in MorphoJ [42] to account for the size differences that are expected to vary between developmental time points. A Procrustes superimposition, which transforms shapes so that they are in maximal superimposition, was applied to the zFACE data in an attempt to remove time dependent variation due to size, position, or rotation and the PCA was repeated; the first 4 PCs now cumulatively explained 92% of the variance (the first 2 PCs explained 84%) (**Supplemental Figure 1A**). Examination of the PC plot for the first two components showed very similar results to the PCA using untransformed data, with developmental days varying across PC1 and no strain-based clusters in the data (**Supplemental Figure 1A**). To focus on how facial shape changed as development progressed, discriminant function analysis (DFA) was utilized and results were summarized by overlaid wireframe representations (**Supplemental Figure 1B-D**). This analysis revealed significant shape changes between 3 and 4 dpf (Procrustes distance = 0.14, p < 0.0001) and 4 and 5 dpf (Procrustes distance = 0.12, p = 0.005), but no changes between 5 and 6 dpf (Procrustes distance = 0.05, p = 0.04 before Bonferroni correction) (**Supplemental Figure 1B-D**). No strain-specific shape differences were observed at any of the developmental time points (**Supplemental Figure 1E-G**). Together, results from these analyses suggest that zFACE represents a robust and sensitive approach for morphometric analysis of facial development in zebrafish embryos.

### Application of zFACE to *smarca4a* mutant zebrafish reveals similarities with Coffin-Siris Syndrome (CSS)

To test the application of zFACE and assess its utility for detection of variation in morphology that occurs after genetic perturbation, we analyzed zebrafish with loss of *smarca4a*, which have been described to have craniofacial anomalies [39,40,41]. Our confocal rostral captures revealed a severely affected facial phenotype in *smarca4a* homozygous mutant larvae at 5 dpf, while heterozygotes and wild-type larvae showed normal developmental hallmarks (**Figure 4A, Supplemental Figure 2**). *Smarca4a* mutants had a narrow head and face, smaller brain, reduced olfactory pits, open and elongated oral cavities and small mandibles (**Figure 4A**). Automated calculation and comparison of zFACE measurements showed that homozygous mutants significantly differed from heterozygotes and wild-type controls in 33 out of the 39 zFACE measurements, including reduced facial width, increased facial height, decreased olfactory distance, reduced mouth width and altered oral cavity angles (ANOVA, post-hoc Tukey’s test, p < 0.0013 for all; **Table 2**). All three groups were equal in upper lip width, chin width, mid olfactory to chin height, mouth area and the difference between chelion and labialie inferius left and right angles, suggesting that *smarca4a* does not affect these structures.

**Figure 4.**
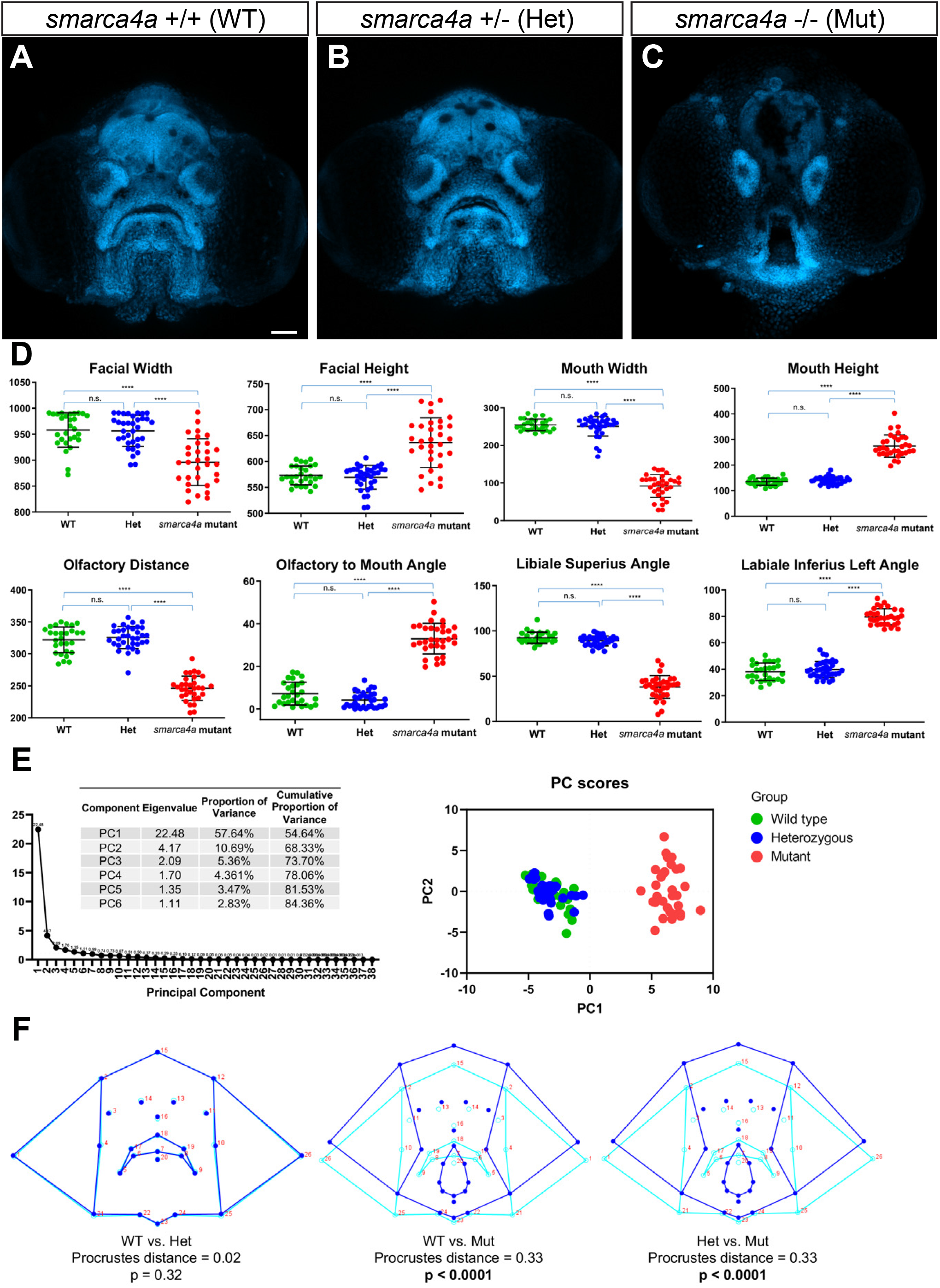
zFACE analysis of zebrafish with loss of *smarca4a*. **A-C.** Confocal images of wild type, WT (**A**), heterozygous, Het (**B**) and homozygous mutant *smarca4a* (**C**) larvae. **D**. zFACE analysis pinpointed 33 significantly altered facial measurements between the groups, with examples of dimensions that were unaltered, decreased and increased shown. **E**. Untransformed zFACE features were subjected to multivariate statistical testing using principal component analysis (PCA). The first 2 components explained 68% of the variance in the dataset and the PC plot showed clear separation of the homozygous mutant group from the WT and heterozygote groups. **F**. Discriminant function analysis (DFA) after landmark data was Procrustes transformed further supported that *smarca4a* homozygous mutants had different average facial shapes compared to either WT or heterozygous larvae, while WT and heterozygous groups had the same facial shape. Wireframe representations of facial shapes highlight the alterations to landmarks in the upper face, eyes, oral cavity, midface and lower jaw in *smarca4a* mutants. Scale bars = 50μm.

**Table 2:** Results of zFACE feature comparison between WT, heterozygous and homozygous mutant *smarca4a* zebrafish embryos.

| **zFACE feature** | **Change in *smarca4a* mutant** | **Het vs. *smarca4a* mutant** |
| --- | --- | --- |
| Height | ↑ | <0.0001 |
| Olfactory to Mouth | ↑ | <0.0001 |
| Olfactory to Mouth 2 | ↑ | <0.0001 |
| Difference | ↑ | <0.0001 |
| Alternate Height | ↑ | <0.0001 |
| Mouth Height | ↑ | <0.0001 |
| Neuromast Angle 1 | ↑ | <0.0001 |
| Neuromast Angle 2 | ↑ | <0.0001 |
| Difference | ↑ | 0.0001 |
| Neuromast Height | ↑ | <0.0001 |
| Area Top | ↑ | <0.0001 |
| Chelion Left Angle | ↑ | <0.0001 |
| Chelion Right Angle | ↑ | <0.0001 |
| Crista Philtri Left Angle | ↑ | <0.0001 |
| Crista Philtri Right Angle | ↑ | <0.0001 |
| Crista Philtri Diff | ↑ | <0.0001 |
| Labiale Inferius Left Angle | ↑ | <0.0001 |
| Labiale Inferius Right Angle | ↑ | <0.0001 |
| Width | ↓ | <0.0001 |
| Olfactory Distance | ↓ | <0.0001 |
| Lower Lip Width | ↓ | 0.0005 |
| Mouth Width | ↓ | <0.0001 |
| Olfactory to Mouth 3 | ↓ | <0.0001 |
| Mouth to Chin | ↓ | <0.0001 |
| Neuromast Width | ↓ | <0.0001 |
| Mid Neuromast Width | ↓ | <0.0001 |
| Average Length Olfactory to mouth | ↓ | <0.0001 |
| Area Bottom | ↓ | <0.0001 |
| Area Combined | ↓ | <0.0001 |
| Mouth Perimeter | ↓ | <0.0001 |
| Libiale Superius Angle | ↓ | <0.0001 |
| Labiale Inferius Angle | ↓ | <0.0001 |
| Labiale Superius Mid Angle | ↓ | <0.0001 |
| Mid Olfactory to Chin height | same | n.s. |
| Chelion Diff | same | n.s. |
| Labiale Inferius Diff | same | n.s. |
| Upper Lip Width | same | n.s. |
| Chin Width | same | n.s. |
| Mouth Area | same | n.s. |

Dimensionality reduction via multivariate PCA of zFACE measurements identified 6 principal components that met the Kaiser cutoff and cumulatively explained 84% of the variance (**Figure 4B**). The first component (PC1) explained 58% of the variance, PC2 explained an additional 11% and PC3 explained 5%. After varimax rotation, component loadings showed that the measurements neuromast height and area top significantly loaded into PC1, while width, neuromast width, average length of olfactory to mouth and facial area combined, loaded into PC2. The scoreplot (PC plot) for these first two components showed *smarca4a* homozygous mutants having higher PC1 scores and separating from wild-type and heterozygous groups (**Figure 4C)**. Using the PCA model, PC scores were predicted for each embryo, and logistic regression was performed with these predicted scores using group as the dependent variable and PC score as the independent variable. Importantly, PC1 score could alone predict whether an embryo was a homozygous *smarca4a* mutant (when PC1 score was greater than −0.98 homozygote mutant genotype was predicted accurately) but could not distinguish between wild-type or heterozygotes (p = 0.86) (**Figure 4C**). Because the zFACE measurements are composed of different types of units (angles, areas, distances and differences) and the standard deviation (SD) between the measurements is not equal, PCA was also performed after standardizing the data and scaling it to have a mean of 0 and SD of 1. This led to very similar PCA results, with 6 PCs cumulatively explaining 88% of the variability. Additionally, we compared 4dpf and 6 dpf wild type embryos (1 day earlier and one day later) to the *smarca4a* embryos to examine if the morphological differences in the mutants could be due to delayed facial development. In the PCA plot, all wild type embryos plotted separately from the *smarca4a* homozygous mutants, suggesting that development is not simply delayed and the abnormal facial phenotype is specific to *smarca4a* disruption (**Supplemental Figure 3**).

Lastly, we performed Procrustes superimposition of the landmark coordinates in MorphoJ [42]. Data was analyzed based on the assumption of object symmetry in the head, and the symmetric portion was evaluated in all analyses [43]. PCA resulted in PC1 accounting for 85% of the variance, PC2 4% and PC3 another 3%; cumulatively these first 3 PCs explained 92% of the variance (**Supplemental Figure 4A**). Similar to results from the untransformed PCA scoreplot, the genotype groups could be clearly distinguished by graphing PC1 vs PC2 (**Supplemental Figure 4B**). Shape changes across PC1 affected mouth sphericity, with landmarks around the oral cavity showing the biggest changes (highest eigenvectors), while changes across PC2 are suggestive of a narrower midface, elongated head and mouth (**Supplemental Figure 4C**).

To compare the 3 genotype groups in an unbiased manner, we used canonical variate analysis (CVA) as an exploratory method. The results showed significant Procrustes distance differences among groups, with the *smarca4a* homozygotes significantly differing from the heterozygote and wild type groups (distance = 0.33 and p<0.0001 when homozygotes were compared to WT and distance = 0.33, p <0.0001 when homozygotes were compared to heterozygotes). Changes in canonical variate 1 (CV1) involved landmarks around the mouth led to a more round and open mouth, while those along CV2 were associated with a narrower and elongated face, similar to the results obtained from the PCA analysis (**Supplemental Figure 4**). Because the WT and heterozygous groups were so similar, discriminant function analysis (DFA), which is similar to CVA but compares only two groups at a time, was performed to focus on the shape changes specific to the homozygous mutants. Results showed a Procrustes distance of 0.33 (p < 0.0001) in both the WT or heterozygous comparison to homozygous mutants (**Figure 4C**), and wireframe representations depicted shape change involving a vertically elongated mouth (**Figure 4D**), reflecting the phenotype seen in the confocal images (**Figure 4A and Supplemental Figure 2**).

To better understand the observed phenotypic abnormalities, we performed fluorescent *in situ* hybridization to examine mRNA expression of *smarca4a* in the craniofacial region. Expression of *smarca4a* mRNA was observed in several facial tissues at 5 dpf, including the perioral and oral tissues, olfactory placodes and the telencephalon in both wild type and heterozygous larvae (**Figure 5**). *Smarca4a-/-* mutants on the other hand, showed a dramatically reduced signal in the entire craniofacial region, suggesting that the point mutation (which creates a premature stop codon) leads to no detectable mRNA expression in developing facial structures (**Figure 5**) [40]. This expression pattern of *smarca4a* mRNA was consistent with the facial structures that zFACE identified as the most altered in the *smarca4a* mutants. Based on the observed morphological changes in brain and neural tissues, width and length of the brain were measured and mutants showed an increased length to width ratio (ANOVA, p < 0.0001) (**Supplemental Figure 5**). Together, these results suggest similarities between *smarca4a* mutants and individuals with Coffin-Siris syndrome (CSS), who often present with craniofacial abnormalities such as a smaller mouth, thicker lips, and reduced philtrum as well as intellectual disability and microcephaly [44,45].

**Figure 5.**
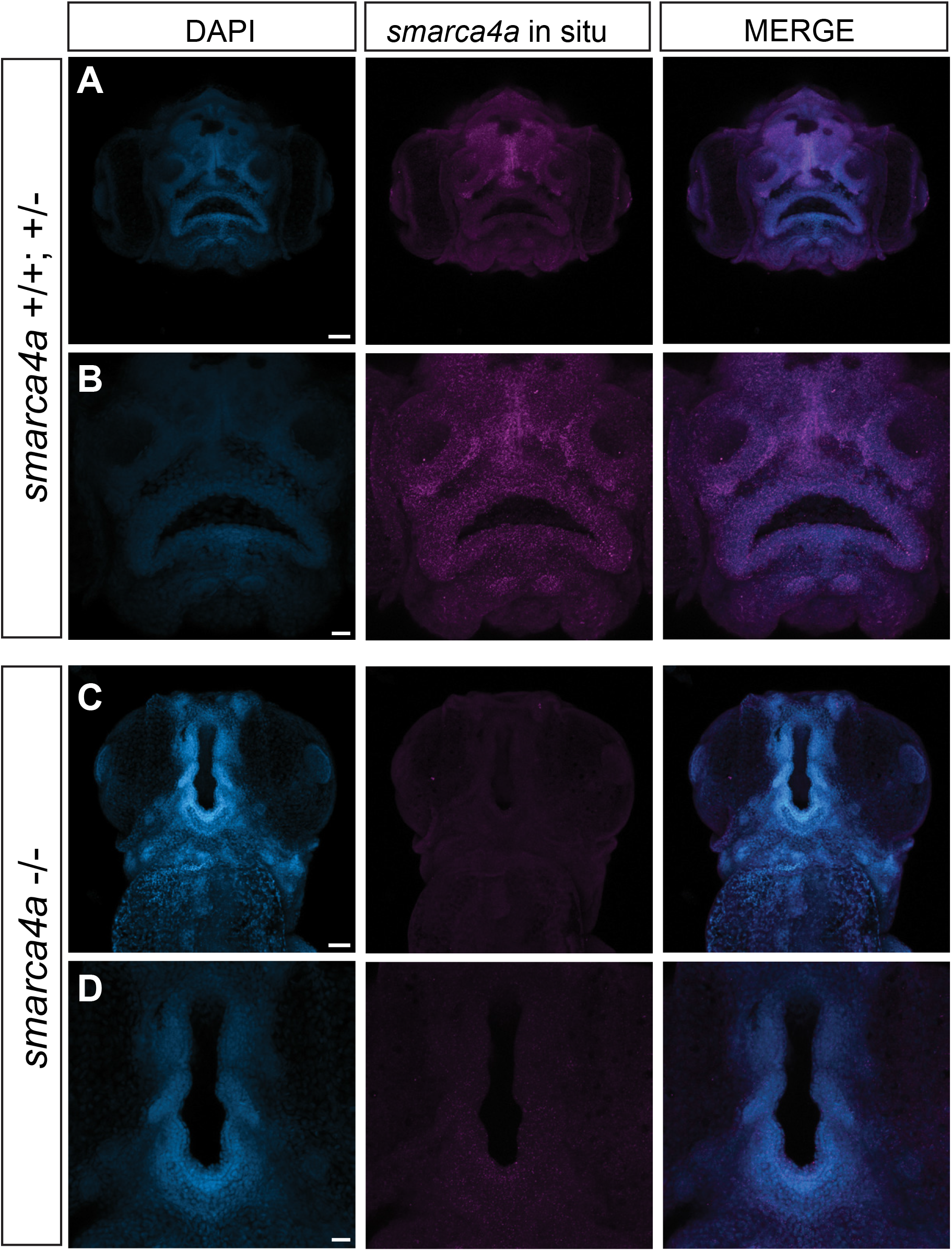
*smarca4a* mRNA expression at 5 dpf. **A-D.** Fluorescent i*n situ* hybridization showed that *smarca4a* is expressed in multiple facial tissues and underlying brain structures in wild type and heterozygous zebrafish (**A-B**), while *smarca4a-/-* mutants showed dramatically reduced mRNA expression (**C-D**). Scale bars in A,C = 50μm, B,D = 20μm.

## DISCUSSION

Geometric morphometrics (GMM) is a powerful quantitative approach for assessing phenotypic differences arising from alterations in shape and size. Here we have developed a streamlined method called zFACE to visualize craniofacial structures and apply GMM to evaluate the developing zebrafish orofacial region. While similar methods exist for other model organisms and human studies, there have been few applications for zebrafish, and zFACE fills this gap in a way that builds on existing approaches and facilitates cross-species comparisons. We first used zFACE to characterize and understand changes in facial morphology from days 3 to 6 of zebrafish development and established standards at various time points. Additionally, we applied it to analyze *smarca4a (yng)* mutants and uncovered morphometric alterations that coincided with where *smarca4a* was expressed, supporting the use of these mutants to inform on mechanisms driving Coffin-Siris syndrome (CSS). Collectively, the development and testing of zFACE presented here, supports its use as a powerful quantitative tool to uncover previously unappreciated craniofacial alterations in zebrafish genetic models.

To facilitate widespread use for future studies, we added several key features to make zFACE informative, reliable and easy to implement. A detailed protocol is included, along with recommendations for ensuring reproducible capture of structures when imaging and placing landmarks. Template files that automatically calculate measurements, perform basic analyses and plot results from user data are provided (**resource materials**). This feature-focused exploration is useful for quickly identifying regions or structures affected by the experimental variable. For example, reduced size can reflect reduced growth during development and point to possible cell population deficiencies driving effects in a particular region, or globally [36,46]. The ability to perform shape analysis from the same landmarks in MorphoJ [42], a widely used and open-source morphometric program, offers additional advantages: its reliable, user-friendly and has been implemented in craniofacial studies in both zebrafish and humans.

Previous studies have relied on lateral and ventral views of zebrafish embryos and larvae, preventing acquisition of information on important aspects of facial morphology. The staining and mounting technique presented here is able to facilitate acquisition of rostral view images. Using this view, standard anatomical locations and nomenclature facilitate comparison with human, mouse and other model organism studies [36,47,48]. However, zFACE is also amendable for expansion to analyze the lateral and ventral views with the selection of landmarks that capture relevant structures in these views. Embryonic/larval zebrafish offer a unique opportunity for detection of small effects due to high-quality imaging capabilities and large numbers of individuals that can be assessed for each condition, providing high-throughput, adequately powered studies. Additionally, we selected a moderate number of landmarks to allow for exploratory analyses such as CVA, for which the number of individuals (n) in each group needs to be higher than the number of landmarks [28]. We also designated a sufficient proportion of the landmarks around the mouth to capture the complex and dynamic shape this structure has during the course of development.

Here, we applied zFACE to test whether we could detect phenotypic changes after genetic manipulation. Using a mutant with previous reported craniofacial abnormalities but analyzed using only lateral and ventral views, our analysis identified several altered facial features in *smarca4a* mutant larvae that contribute to a different overall average face shape compared to wild type and heterozygous animals. The abnormal facial features, together with the alterations in brain morphology, show parallels to those observed in patients with Coffin-Siris syndrome (CSS). CSS is a rare congenital disorder that presents with distinct facial features due to craniofacial abnormalities, fifth digit anomalies, microcephaly and intellectual disability [49,50]. The clinical features are heterogenous and variants in several genes encoding proteins in the SWI/SNF complex, including *SMARCA4A*, have been identified as causal, with variants spanning the gene and disrupting several protein domains [49,51]. To date, there are no established animal models for the subtype of CSS with *SMARCA4A* mutations [52,53]. Thus, the *smarca4a* zebrafish larvae provide an opportunity for future studies to better understand disruptions in development and identify potential strategies for therapeutic intervention.

The zebrafish *smarca4a* (also *yng* or *brg1*) mutant, has a point mutation (C to A transversion) leading to a premature stop codon early in the gene, preceding all functional domains [40]. This mutant has been described as having craniofacial abnormalities, alterations in brain size and patterning and other anomalies in neural crest cell derived tissues [39]. The current study provides high-resolution phenotypic information to further support these abnormal craniofacial characteristics of *smarca4a* mutants compared to wild type and heterozygous controls. Our results identified important similarities in craniofacial features of the mutant with abnormalities reported for CSS. Our results also found severely abnormal oral cavities, suggesting compromised oropharynx function and ability to feed, and is possibly another reason why these mutants only survive to 7 days post fertilization [41]. The frontal view additionally allowed brain measurements of the telencephalon, which showed an altered length to width axis, supporting that mutant *smarca4a* leads to abnormal brain morphology [39]. Future studies can take a similar approach to integrate phenotypic data with genetic and molecular information for greater mechanistic insights on gene functions during development.

The complexity of craniofacial development and the structures that make up the craniofacial regions make the detection of subtle phenotypic variation challenging and require advanced analytical tools. zFACE can be implemented in a way that’s sufficiently powered to detect subtle morphological changes and better allow for genotype-phenotype correlations, which can serve an important role in the investigation of multifactorial disorders, gene-environment interactions, pharmacological interventions, and other studies. Direct comparisons on how morphology changes based on genetic perturbations can also bridge the gap between results from GWAS studies and the biological effects of genetic variation on craniofacial structures. In conclusion, zFACE can facilitate important studies to examine and integrate morphological effects of genetic, environmental or developmental perturbations in zebrafish studies of craniofacial development.

## ACKNOWLEDGEMENTS

This work was supported by the Gulf Coast Consortium (GCC) Dunn Collaborative Research Award to GTE and JTH; the National Institute of Dental and Craniofacial Research R01DE011931 to JTH; and the Cancer Prevention Institute of Texas, RR14007, National Institutes of General Medical Sciences, R01GM124043, and the Linda and Mark Quick Award for Basic Science to GTE.

## AUTHOR CONTRIBUTIONS

S.T.L., O. E. R. and G.T.E conceived the methodology and analysis pipeline. L.M. enhanced the zFACE methodology with advanced statistics and shape analysis. L.M., P.K., S.T.W. and O.E.R. performed all the experiments and subsequent analysis. S.S.H. provided assistance with statistical analysis. J.T.H. provided support and insight to optimize the methodology. L.M. and G.T.E. wrote the manuscript.

## METHODS

### Experimental Model and Subject Details

Experiments were conducted on larval zebrafish (*Danio rerio*) maintained under standard laboratory conditions with a cycle of 14 h of light and 10 h of darkness. Larvae were collected and kept in E3 larva medium at 28.5°C and staged as previously described [54]. The zebrafish used in this study were handled in accordance with the guidelines of the University of Texas MD Anderson Cancer Center Institutional Animal Care and Use Committee and UTHealth Animal Welfare Committee.

*Smarca4a ^a8^ (yng)* mutant zebrafish [40] were obtained from the Zebrafish International Resource Center (ZIRC).

### Specimen collection, staining, mounting and image acquisition

Specimens were collected and fixed overnight on a shaker in 4% fixation solution prepared from dilution of a 36.5 % formaldehyde stock solution (Formalin; Simga) into PBS with a small amount of detergent (0.05% Triton-X 100) (0.05% PBST).

Fixation solution was then removed, briefly washed with PBST and DAPI (4’,6-diamidino-2-phenylindole, Thermo Scientific) was added to PBST at a 1 in 1000 dilution. The samples were incubated in the DAPI solution for 30 minutes and rinsed with 0.05% PBST in a series of three 20 minute washes. Samples were then rinsed and stored in PBS at 4° C. Each embryo was mounted in a 35 mm glass bottom culture dish with a 10mm microwell (MatTek Corporation) filled with 1% low-melt agarose solution. For the frontal/rostral orientation, each embryo was manipulated by moving the tail to suspend the sample upside down, ensuring that the eyes were on the same plane to reduce variability in mounting angles.

To ensure proper orientation of each sample, the clear visualization of the midline between the brain ventricles and the ability to see folds in lower jaw tissues was used as a guide. If the size of the embryo head was too big, the stage was rotated 45 degrees during image acquisition so both eye lenses could be captured and the last Z-stack ensured inclusion of these structures. Typically, the 10x (magnification) objective was used to capture 40 slices of 3.08 uM thickness at 2% laser power using 600V, digital offset of 2 and digital gain of 1. The images were 1024 x1024 at 8bit/pixel and the maximum scan speed and averaging of 2 was used.

### Coordinate Point System

A 26-point system was utilized for landmarks in the craniofacial region of zebrafish larvae. Each point was assigned a number 1 through 26. Confocal images were opened in ImageJ, points were selected in numerical order using the Point Picker tool, and X Y coordinates for each point were extracted.

### Facial dimension measurements

Using these X Y coordinates and the distance formula (√(*x*_2_ – *x*_1_)^2^ + (*y*_2_ – *y*_1_)^2^), various distance, angular, and area measurements were calculated between landmarks. Angular measurements utilized the formula *A*^2^ = *B*^2^ + *C*^2^ – 2*BC* cos *a*, where *a* is the angle of the BC connecting point. Triangular areas were found using 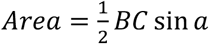. All calculations were performed using Microsoft Excel.

### Statistical analysis of zFACE measurements

For statistical analysis of each zFACE measurement, the GraphPad Prism analysis function was used to run one-way ANOVA with Tukey’s post hoc tests for multiple comparisons. Bonferroni correction for 39 tests was applied and p < 0.00128 was considered significant.

Multivariate analysis of the zFACE measurements was performed in Stata and GraphPad Prism. Data was standardized and principal component analysis (PCA) was performed. Multiple models were run to thoroughly identify/evaluate the most robust model in Stata, where data was rotated and PC loadings could be calculated. Once modeling was determined in Stata, a rapid and streamlined analysis was rerun in Graphad Prism, producing similar results.

### Shape analysis of zFACE landmark coordinates

Landmark coordinates were imported into MorphoJ and standard protocols were utilized to perform Procrustes superimposition on principal axes as well as principal component analysis (PCA), canonical variate analysis (CVA) and discriminant function analysis (DFA) to compare shape changes across the dataset and between groups [42].

### Genotyping of *smarca4a* mutants

*Smarca4a ^a8^ (yng)* mutant zebrafish [40] were identified by a PCR reaction using the following primer sequences: Forward 5’-CCT GTC ATG CCC CCT CAG AC-3’; and Reverse 5’-CCG ACC CCC ACT TTG AGA AC-3’. The resulting 190base pair band was excised and a restriction digest using RsaI was performed at 37C for four hours. RsaI only cuts the WT band, therefore, this results in wild-type band having 2 bands (50 + 140 bp), heterozygous larvae with 3 bands (50 + 140 +190 bp) and homozygous mutants with one 190bp band.

