## Supplemental Data for "zFACE: <u>F</u>acial <u>A</u>nalytics from a <u>C</u>oordinate <u>E</u>xtrapolation System for Developing Zebrafish"

#### SUPPLEMENTAL FIGURE LEGENDS

**Supplemental Figure 1. Shape analysis using zFACE landmark coordinates. A.** PCA was performed after Procrustes superimposition of the zFACE landmark coordinates. The first 2 components explained 84% of the variance in the dataset, and PC plots showed similar clustering of data by developmental day across PC1, while no strain-specific clustering was observed. **B-D.** Discriminant function analysis (DFA) was utilized to identify and follow shape differences between developmental days and between strains. Significant shape changes were found between 3 and 4 dpf and 4 and 5 dpf, while face shape was not different between 5 and 6 dpf. **E-F.** There were no strain-specific differences.

**Supplemental Figure 2. Phenotype of *smarca4a* mutant larvae**. Facial phenotypes for wild-type, heterozygous and *smarca4a* homozygous mutant larvae at 5dpf.

**Supplemental figure 3: Comparison of *smarca4a* mutants with other developmental timepoints.** PCA model showing that the facial phenotype of *smarca4a* homozygous mutants at 5dpf is different from 4 and 6dpf wild-type larvae.

**Supplemental Figure 4. Analysis of *smarca4a* mutant zebrafish larvae. A.** PCA results after Procrustes superimposition. **B.** The component plot with confidence ellipses shows overlap of the wild type and heterozygous *smarca4a* larvae and clear separation of the homozygous mutants from the other two groups. **C.** Transformation grids with lollipop graphs showing the resulting facial shape changes in DFA of *smarca4a* wild type, heterozygous and homozygous mutant larvae.

**Supplemental Figure 5. Altered brain morphology in *smarca4a* mutants.** Brain length and width were measured for all genotype groups on the rostral confocal images. S*marca4a* homozygous mutants showed increased length to width ratio compared to wild type and heterozygote larvae, indicating morphological changes in the telencephalon. **** $p < 0.0001$

**Supplemental Table 1. zFACE landmark and measurement definitions.**

**Supplemental Table 2. Component loadings of zFACE measurements after promax rotation for PCA analysis of development.**

### Supplemental Figure 1.

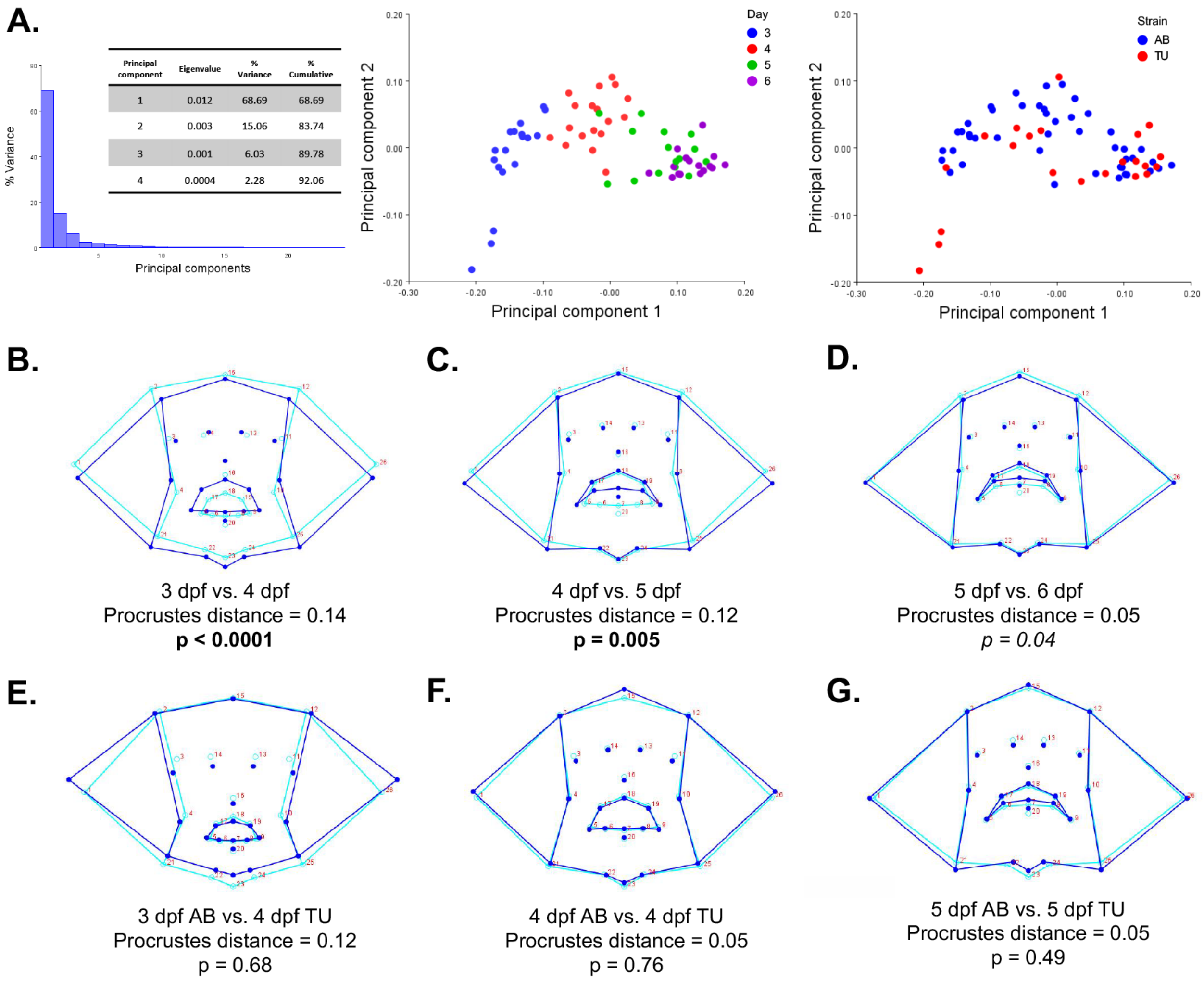


| Principal component | Eigenvalue | % Variance | % Cumulative |
|---|---|---|---|
| 1 | 0.012 | 68.69 | 68.69 |
| 2 | 0.003 | 15.06 | 83.74 |
| 3 | 0.001 | 6.03 | 89.78 |
| 4 | 0.0004 | 2.28 | 92.06 |

B. 3 dpf vs. 4 dpf
Procrustes distance = 0.14
**p < 0.0001**

C. 4 dpf vs. 5 dpf
Procrustes distance = 0.12
**p = 0.005**

D. 5 dpf vs. 6 dpf
Procrustes distance = 0.05
*p = 0.04*

E. 3 dpf AB vs. 4 dpf TU
Procrustes distance = 0.12
p = 0.68

F. 4 dpf AB vs. 4 dpf TU
Procrustes distance = 0.05
p = 0.76

G. 5 dpf AB vs. 5 dpf TU
Procrustes distance = 0.05
p = 0.49

**Supplemental Figure 2.**

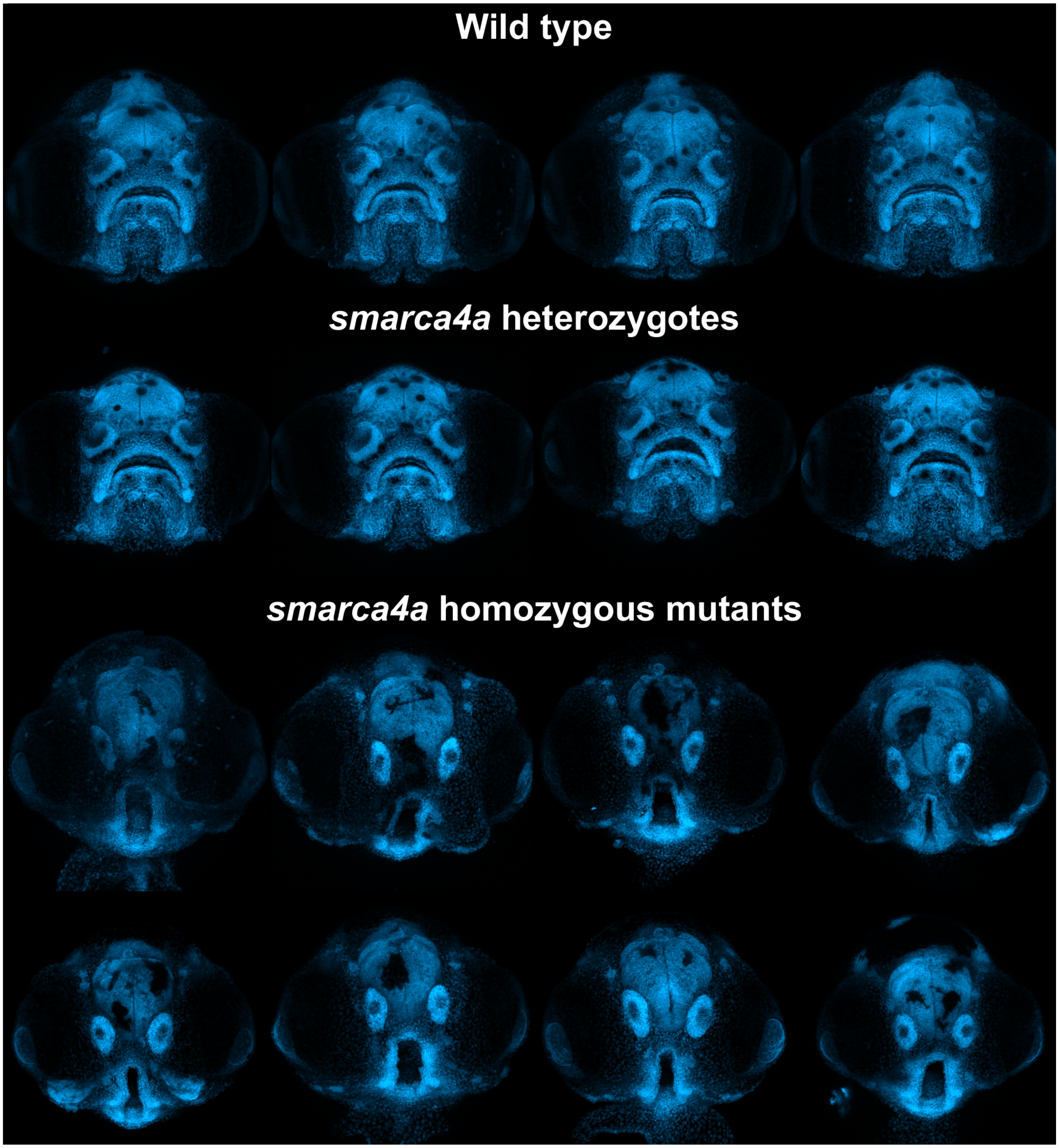

**Supplemental Figure 3.**

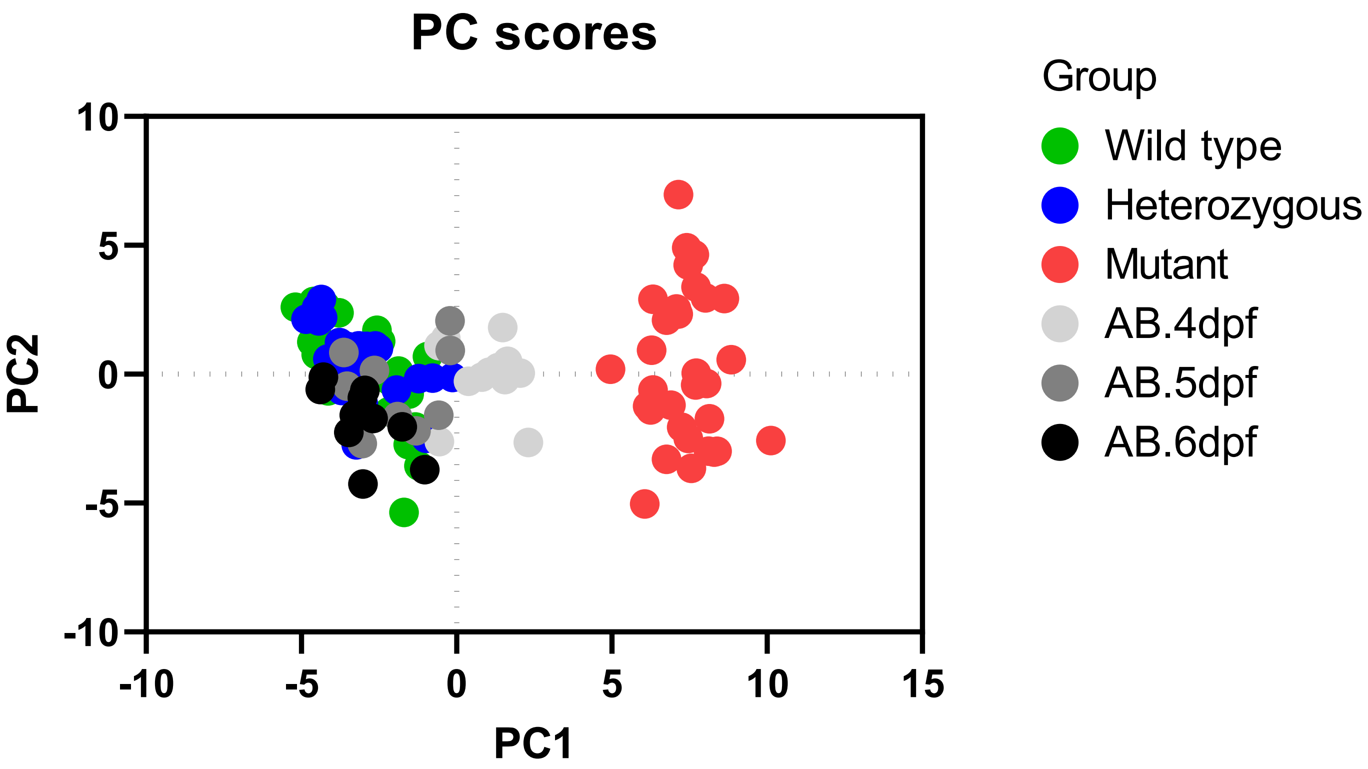

### Supplemental Figure 4.

**A.**

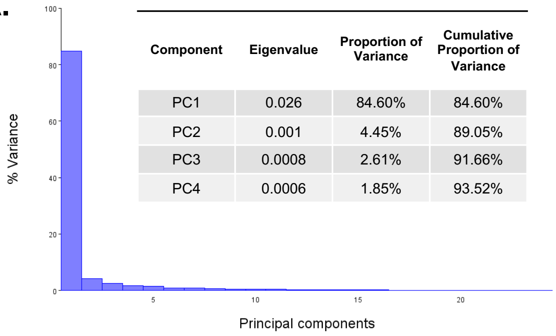


| Component | Eigenvalue | Proportion of Variance | Cumulative Proportion of Variance |
|---|---|---|---|
| PC1 | 0.026 | 84.60% | 84.60% |
| PC2 | 0.001 | 4.45% | 89.05% |
| PC3 | 0.0008 | 2.61% | 91.66% |
| PC4 | 0.0006 | 1.85% | 93.52% |

**B.**

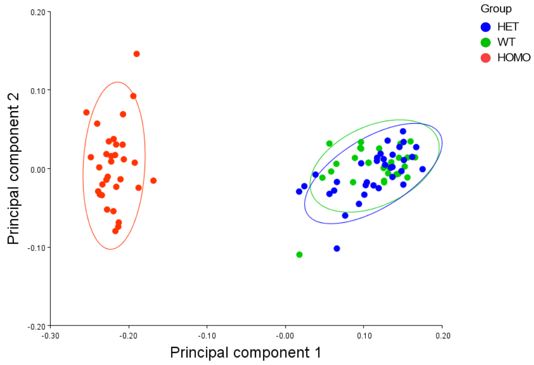


**C.**

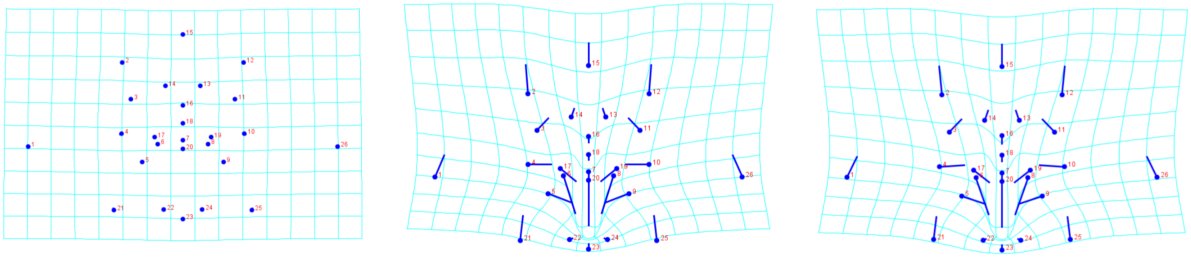

WT vs. HET    WT vs. HOMO    HET vs. HOMO

### Supplemental Figure 5.

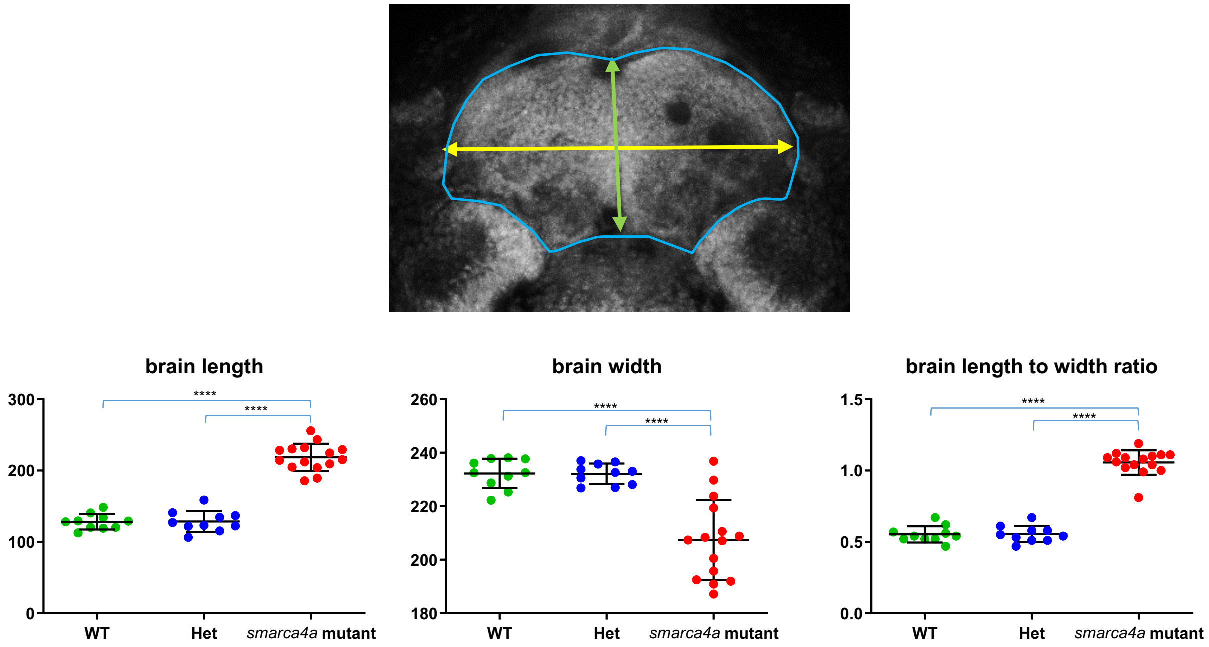

### Supplemental Table 1.

Landmarks:

| Landmark | Name | Definition |
|---|---|---|
| **1** | Right eye pupil | Right pupil apex point |
| **2** | Right dorsal neuromast | Center of the right neuromast on top of the olfactory placode |
| **3** | Right olfactory placode | Point in the center of the olfactory placode |
| **4** | Right middle neuromast | Center of the right neuromast below the olfactory placode |
| **5** | Right chelion | Point at the junction of the upper lip and lower lip, located on the right labial commissure |
| **6** | Midpoint from right chelion to labiale inferius | Midpoint from right chelion to labiale inferius |
| **7** | Labiale inferius | Midpoint of the lower vermillion line |
| **8** | Midpoint from left chelion to labiale inferius | Midpoint from left chelion to labiale inferius |
| **9** | Left chelion | Point at the junction of the upper lip and lower lip, located on the left labial commissure |
| **10** | Left middle neuromast | Center of left neuromast below the olfactory placode |
| **11** | Left olfactory placode | Point at the center of the left olfactory placode |
| **12** | Left dorsal neuromast | Center of the left neuromast on top of the olfactory placode |
| **13** | Left midface neuromast | Center of the left neuromast between the olfactory placodes |
| **14** | Right midface neuromast | Center of the right neuromast between the olfactory placodes |
| **15** | Dorsal midpoint of face | Located in the interhemispheric brain region |
| **16** | Supralabiale | Center of the upper border of the upper lip |
| **17** | Right christa philtri | The point on each elevated margin of the philtrum just above the vermilion line |
| **18** | Labiale superius | Midpoint on the lower border of the upper lip |
| **19** | Left christa philtri | The point on each elevated margin of the philtrum just above the vermilion line |
| **20** | Sublabiale | Center of the lower border of the lower lip |
| **21** | Right ventral neuromast | Center of the right ventral neuromast |
| **22** | Indentation on right of gnathion | Point where the gnathion meet on the right |
| **23** | Gnathion (menton) | Lowest median landmark on the lower border of the mandible (Menton: most inferior median point of the chin) |
| **24** | Indentation on left of gnathion | Point where the gnathion folds meet on the left |
| **25** | Left ventral neuromast | Center of the left ventral neuromast |
| **26** | Left eye pupil | Left pupil apex point |

Measurements:

| Measurement | Name | Definition |
|---|---|---|
| **1** | Width | Point Measure from point 1 to 26 |
| **2** | Height | Measurement from point 15 to 23 |
| **3** | Olfactory distance | Measurement from point 3 to 11 |
| **4** | Upper lip width | Measurement from point 16 to point 18 |
| **5** | Lower lip width | Measurement from point 7 to point 20 |
| **6** | Mouth width | Measurement from point 5 to point 9 |
| **7** | Olfactory to mouth | Angle 11 to 3 to 16 |
| **8** | Olfactory to mouth 2 | Angle 3 to 11 to 16 |
| **9** | Olfactory difference | Difference of Olfactory to Mouth 1 and Olfactory to Mouth 2 |
| **10** | Olfactory to mouth 3 | Angle 3 to 16 to 11 |
| **11** | Chin width | Measurement from point 21 to point 25 |
| **12** | Mouth to chin | Measurement from point 16 to point 23 |
| **13** | Alternate height | Midpoint from points 2 to 12 to point 23 |
| **14** | Mouth height | Measurement from point 16 to point 20 |
| **15** | Neuromast angle 1 | Angle 2 to 4 to 16 |
| **16** | Neuromast angle 2 | Angle 12 to 10 to 16 |
| **17** | Neuromast difference | Difference between neuromast angle 1 and neuromast angle 2 |
| **18** | Neuromast height | Midpoint from points 2 to 12 to midpoint from points 4 to 10 |
| **19** | Neuromast width | Midpoint from points 2 to 4 to midpoint from points 12 to 10 |
| **20** | Mid neuromast width | Measurement from point 13 to point 14 |
| **21** | Average length olfactory to mouth | Average measurement from points 3 to 16 and from points 11 to 16 |
| **22** | Area top | Area from measurement of points 2 to 12 to 10 to 4 |
| **23** | Area bottom | Area from measurement of points 4 to 10 to 25 to 21 |
| **24** | Area combined | Sum of value from area top and area bottom |
| **25** | Mouth area | Sum of areas from points 5 to 17 to 18 and points 9 to 19 to 18 |
| **26** | Mouth perimeter | Sum of measurements from points 5 to 17, 18 to 17, 19 to 18, 9 to 19, 8 to 9, 7 to 8, 6 to 7, and 5 to 6 |
| **27** | Mid olfactory to chin height | Midpoint from points 3 to 11 to point 23 |
| **28** | Labiale superius angle | Angle 5 to 18 to 9 |
| **29** | Chelion left angle | Angle 5 to 9 to 18 |
| **30** | Chelion right angle | Angle 9 to 5 to 18 |
| **31** | Chelion difference | Difference between the left and right chelion angles |

| | | |
|---|---|---|
| **32** | Labiale inferius angle | Angle 17 to 7 to 19 |
| **33** | Christa philtri left angle | Angle 17 to 19 to 7 |
| **34** | Christa philtri right angle | Angle 19 to 17 to 7 |
| **35** | Christa philtri difference | Difference between left and right christa philtri angles |
| **36** | Labiale superius mid angle | Angle 6 to 18 to 8 |
| **37** | Labiale inferius left angle | Angle 6 to 8 to 18 |
| **38** | Labiale inferius right angle | Angle 8 to 6 to 18 |
| **39** | Labiale inferius difference | Difference between the Labiale Inferius Left Angle and Labiale Inferius Right Angle |

#### Supplemental Table 2.

| Variable | PC1 | PC2 | PC3 | PC4 | PC5 | PC6 | Unexplained |
|---|---|---|---|---|---|---|---|
| Width | | | | | | | 0.11 |
| Height | | | | | | | 0.14 |
| Olfactory Distance | | | | 0.51 | | | 0.12 |
| Upper Lip Width | | | | | | -0.38 | 0.37 |
| Lower Lip Width | | | | | | | 0.53 |
| Mouth Width | | | | | | | 0.07 |
| Olfactory to Mouth | | | | | | | 0.04 |
| Olfactory to Mouth 2 | | | | | | | 0.05 |
| Difference | | | | | 0.30 | | 0.48 |
| Olfactory to Mouth 3 | | | | | | | 0.04 |
| Chin Width | | | | | | | 0.15 |
| Mouth to Chin | | | | | | | 0.03 |
| Alternate Height | | | | | | | 0.10 |
| **Mouth Height** | | 0.41 | | | | | 0.05 |
| Neuromast Angle 1 | | | | | | | 0.11 |
| Neuromast Angle 2 | | | | | | | 0.16 |
| Difference | | | | | | 0.77 | 0.25 |
| Neuromast Height | | | | | | | 0.10 |
| **Neuromast Width** | 0.32 | | | | | | 0.10 |
| Mid Neuromast Width | | | | 0.38 | | | 0.24 |
| Average Length Olfactory to mouth | | | | 0.47 | | | 0.05 |
| Area Top | | | | 0.36 | | | 0.10 |
| Area Bottom | | | | | | | 0.04 |
| Area Combined | | | | | | | 0.06 |
| Mid Olfactory to Chin height | | | | | | | 0.10 |
| **Mouth Area** | | 0.32 | | | | | 0.13 |
| Mouth Perimeter | | | | | | | 0.05 |
| Libiale Superius Angle | | | -0.42 | | | | 0.10 |
| Chelion Left Angle | | | 0.45 | | | | 0.13 |
| Chelion Right Angle | | | 0.36 | | | | 0.21 |
| Chelion Diff | | | | | 0.70 | | 0.31 |
| **Labiale Inferius Angle** | | -0.33 | | | | | 0.03 |
| **Crista Philtri Left Angle** | | 0.33 | | | | | 0.03 |
| **Crista Philtri Right Angle** | | 0.32 | | | | | 0.05 |
| Crista Philtri Diff | | | | | | | 0.24 |
| Labiale Superius Mid Angle | | | | | | | 0.02 |
| Labiale Inferius Left Angle | | | | | | | 0.09 |
| Labiale Inferius Right Angle | | | | | | | 0.06 |
| Labiale Inferius Diff | | | | | | | 0.28 |
